# An Information Processing Pattern from Robotics Predicts Unknown Properties of the Human Visual System

**DOI:** 10.1101/2024.06.20.599814

**Authors:** Aravind Battaje, Angelica Godinez, Nina M. Hanning, Martin Rolfs, Oliver Brock

## Abstract

We tested the hypothesis that an algorithmic information processing pattern from robotics, Active InterCONnect (AICON), could serve as a useful representation for exploring human vision. We created AICON-based computational models for two visual illusions: the shape-contingent color aftereffect and silencing by motion. The models reproduced the effects seen in humans and generated surprising and novel predictions that we validated through human psychophysical experiments. Inconsistencies between model predictions and experimental results were resolved through iterative model adjustments. For the shape-contingent color aftereffect, the model predicted and experiments confirmed weaker aftereffects for outline shape manipulations and individual differences in perceived aftereffects. For silencing by motion, the model predicted and experiments validated unexpected trends as well as individual differences. Our findings demonstrate AICON’s ability to capture relevant aspects of human visual information processing including variability across individuals. It highlights the potential for novel collaborations between synthetic and biological disciplines.

## Introduction

We discovered a commonality in illustrations of an algorithm from robot perception and depictions of information processing in the human visual system (see Fig. 1). This tenuous and seemingly far-fetched relationship led us to formulate the central hypothesis of this paper: The algorithmic information processing pattern developed in robotics (AICON: Active InterCONnect) serves as a useful representation for exploring aspects of the human visual system. In this paper, we successfully test this hypothesis in the context of two different visual illusions, the shape-contingent color after-effect [1] and the motion silencing illusion [2]. Although we investigate only in the visual domain, we believe our results may have implications that reach beyond the study of the human visual system.

**Figure 1:**
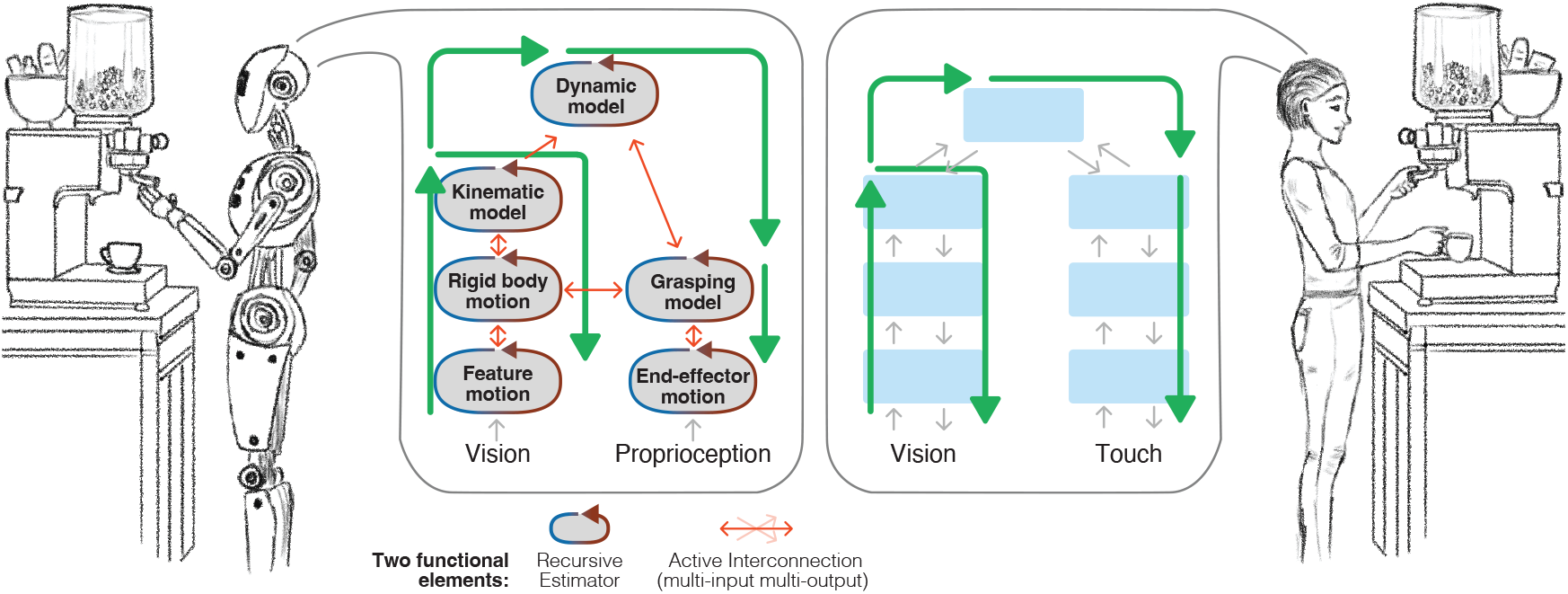
Robotic models for interactive perception, and information processing in humans have striking high-level similarities. (Left) Schematic demonstrating information processing in robots is adapted from Martín-Martín et al. [3, Fig. 9], where blocks correspond to Recursive Estimators and orange arrows correspond to Active Interconnections that show first-order information exchange paths between Recursive Estimators. Recursive Estimators and Active Interconnections are the functional elements of AICON that allow building robust robotic perceptual algorithms. (Right) The high-level information processing architecture in human cortex is adapted from Hawkins et al. [4, p. 118, Fig. 4], where boxes are taken to be functional units in the cortex and gray arrows show first-order information paths. (Both) In both schemes, information travels bottom-up and top-down through different paths as indicated by green arrows, making higher-order information paths. This enables exchange of interdependent information, ultimately allowing Recursive Estimators/functional units to mutually benefit from each other and becoming robust as a whole.

The resemblance in depictions (Fig. 1) may stem from the same essential property: highly interdependent processes with rich interactions. Interdependent processes abound in humans—e.g., linking action and perception [5–7], different sensory modalities [8, 9], or perception and cognition [10, 11]—and may even be a hallmark of intelligence [3, 12]. Interdependent processes are also central to the design of AICON, the information processing pattern from robotics. This makes AICON a promising candidate to analyze properties of human information processing.

To understand how well AICON can be used to explore human vision, we studied two striking visual illusions [1, 2] that demonstrate interactions between shape and color perception [1], and motion-based silencing of perceived changes in stimulus features [2]. For each illusion, we created a computational model using AICON with the goal of reproducing the percepts observed in humans. To establish how well each model generalizes, we obtained predictions for untested conditions and subsequently validated those predictions using human psychophysical studies. When we encountered inconsistencies, we adjusted the models and made further predictions. In this process, we made continuous progress in understanding the perceptual properties underlying the studied phenomena.

Here, we present evidence that AICON— a modeling framework originally developed for robotics—serves as a useful medium to study human vision by accurately capturing relevant aspects of its information processing. Following such an approach opens the door for a novel type of fruitful collaboration between synthetic disciplines and those studying human perception or even behavior, using models directly inspired from robotics and AI. When these models closely capture the regularities of information processing in humans, they allow us to select hypotheses for future experiments that are informed by the algorithmic structure encapsulated by the models. This interdisciplinary approach delivers novel modeling abilities at previously unexplored levels of abstraction, with the potential to substantially accelerate progress in a variety of disciplines studying human perception and behavior.

## Results

### Information Processing Pattern from Robotics (AICON)

Before we detail the studies of human vision we undertake in this work, we must first understand how AICON (Active InterCONnect) is uniquely positioned to help model human perceptual processes.

AICON was shaped by the need to generate robust behavior in real-time from high-dimensional sensory inputs. Fig. 1 (middle-left) shows a schematic for a perceptual model from Martín-Martín et al. [3], where the task was to infer kinematic structure of mechanisms, such as how the coffee filter handle is mechanically connected to the machine, from very high-dimensional visual and proprioceptive inputs. The robot can then use this knowledge to determine how to remove the coffee filter (e.g., turn it counterclockwise).

To run such a task in real-time, the signal processing was factorized into smaller parts, as is common in engineered systems. However, the key insight was recognizing that these factorizations are not independent: the result of signal processing in one part, say Part A, can be used to refine the result in another part, say Part B, and that refined result can then be used to further refine the result in Part A. This co-dependence exists between many parts of the system, allowing information in one part to bootstrap other parts and ultimately enhance the entire perceptual process. Martín-Martín et al. [3] demonstrate that while the factorizations themselves can be relatively straightforward, the effectiveness and robustness of the robotic perceptual system largely depend on the *interactions* that arise from the co-dependence of these *parts*.

To operationalize these insights, AICON provides two functional elements to explicitly express the *parts* and *interactions* in a perceptual system.

1. A **Recursive Estimator** is a *part* that estimates a quantity characterizing a physical process, and it infers this quantity from (possibly) noisy inputs, while taking advantage of temporal structures in the modeled physical process (dynamics). Notable instances of Recursive Estimators include Kalman Filters and Particle Filters [13]. In AICON, these Recursive Estimators are modified to interoperate with Active Interconnections.
2. An **Active Interconnection** encodes the *interactions* between the *parts*. An Active Interconnection connects to several Recursive Estimators and determines how the values of the quantities in Recursive Estimators should be related to each other according to a specified function. It serves as a dynamically adapting medium for the connected Recursive Estimators to adjust themselves and bring each other in agreement.

When multiple Recursive Estimators are connected through appropriate Active Interconnections the interdependent processes bolster each other, and the collective ends up representing dynamically-adjusting complex structures in information, often recovered from high-dimensional raw sensory inputs. From a practitioner’s view, AICON provides an effective way to express robotic perceptual models that are robust to various types of noise and interference. (For further details on AICON, see SI Sec. S1.)

How does AICON help model human information processing? Another way to look at AICON is that it provides a structured way to model highly interdependent processes. As we suppose interdependent processes are common in human information processing [5–10], we expect AICON to provide a useful abstraction to model parts of the human perceptual system. As we show in this paper, this abstraction not only helps create suitable models, but those models also predict certain aspects of human vision. Due to this property, we could quickly co-evolve the models: adjust the models to outcomes of pyschophysical experiments, and design subsequent experiments from the models’ predictions, allowing us to continuously refine our understanding of the studied phenomena. In order, to scrutinize how well AICON captures perceptual properties of human vision, we studied two visual phenomena [1, 2]. We describe our results from these studies in the following.

### Illusion 1: Fill-in Color After-Effect Illusion

In our first study, we turn to the illusion by van Lier et al. [1]. The basic version of the illusion involves adapting to two colored shapes, with one of the shapes partially overlapping the other (Fig. 3a). Subsequently, presenting an outline of one of the shapes elicits an afterimage whose color not only fits the corresponding (negative) color in the adaptation phase, but also fills in the overlapping area (refer to SI Movie S1).

### Illusion From The Model’s Perspective

This illusion shows a close interaction between shape and color perception in human vision. The obvious direction of influence is color-to-shape: color cues delineate different shapes. However, the less obvious direction of influence betrayed by this illusion is shape-to-color: shapes are able to influence the (negative) color you perceive. The model captures this bidirectional nature of influence between two basic perceptual processes including their temporal dynamics and perceptual uncertainty.

Fig. 2a shows a schematic of this model. Here, the input stream is a sequence of images that at first are separated into independent modalities: brightness and color. This information is independently used by shape and color processing that are modeled using Recursive Estimators respectively. As multiple shapes can occupy the same location in the stimulus, the shape processing is modeled to detect and track multiple shapes with a notion of “dis-similarity” to disambiguate between shapes.

**Figure 2:**
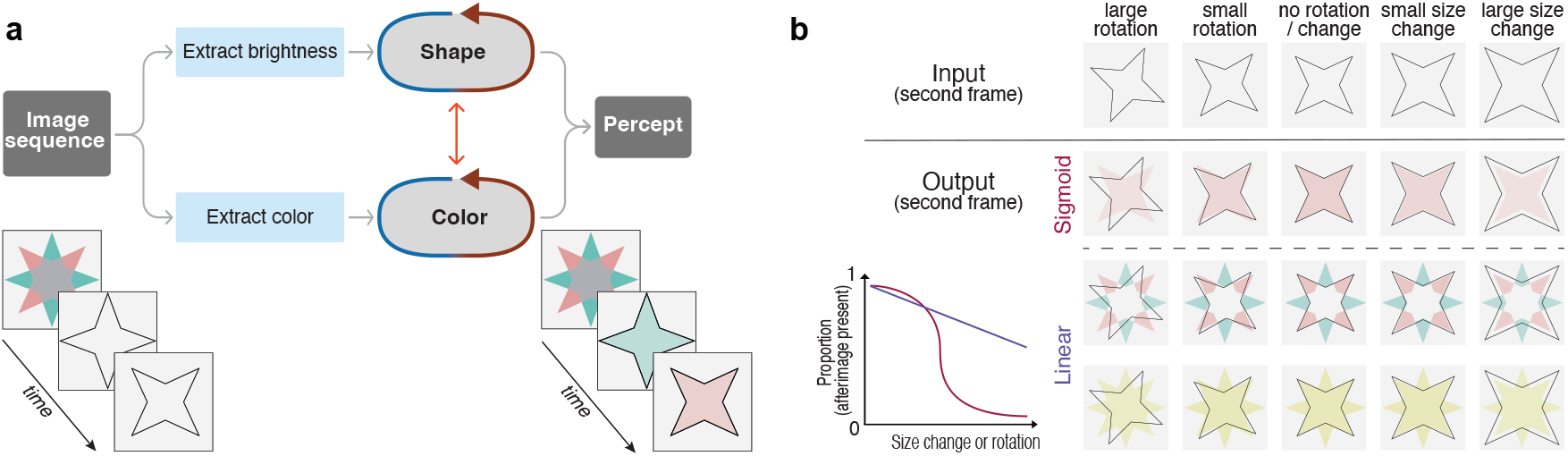
The model produces outputs similar to human percepts, including previously unreported observations. (a) The model encodes the interaction between shape and color perception. Given a sequence of images as input, the model produces images (“percept”) qualitatively similar to that in humans. (b) Different types of aftereffects for different input and model parameter settings: The first row shows various input stimuli variations as performed in Exp. 1A and Exp. 1B. The second row shows corresponding model outputs, where larger shifts in outline relative to the adapting stimulus produce weaker filled-in aftereffects. The third and fourth rows correspond to two different model parameter settings, that were done in order to reconcile with subjects who responded linearly in Exp. 1A. The aftereffects in the third and fourth row correspond to “8-pointed multi-colored star” and “8-pointed single colored star” responses in Exp. 1B, respectively.

In the model, the shape and color processing also exchange overlapping information between them through an Active Interconnection. This Active Interconnection effectively modulates the certainty in each Recursive Estimator depending on the presence of a fitting cue in the complementary estimation process. Finally, the information from Recursive Estimators, including uncertainties associated with the quantities being tracked, are used to interpret the final “percept” for this illusion. (For further details on model, see SI Sec. S3.)

This initial model qualitatively reproduced the effect seen in humans. (The model produces afterimages in positive colors, as opposed to the negative colors in humans. This does not drastically change our inference. More details are provided under Discussion section.) That is, the model produced shape-dependent, filled-in aftereffects with colors that correspond to the initial stimulus, and the strength of aftereffect gradually decreased with time. With this, we made the first model prediction for an untested condition: As a consequence of the shape estimation process being modulated by “dis-similarity”, small modifications to the outline presented after the adapting colored stimulus resulted in weaker aftereffects. Fig. 2b (first row) shows example inputs to the model and Fig. 2b (second row) shows model responses for those stimuli with manipulations. Given these predictions, we conducted psychophysical studies to determine if they hold true in humans.

### Humans Perceive Weaker Aftereffects With Outline Manipulations

In Experiment 1A (Fig. 3a), we investigated the strength of the color aftereffect illusion to delineate the boundaries of coupling between shape and color. We achieved this by altering the size or rotation of the outline relative to the adapting stimulus and then prompting participants to report their perceptions. (Refer to SI Movie S2–S4 for example stimuli.)

**Figure 3:**
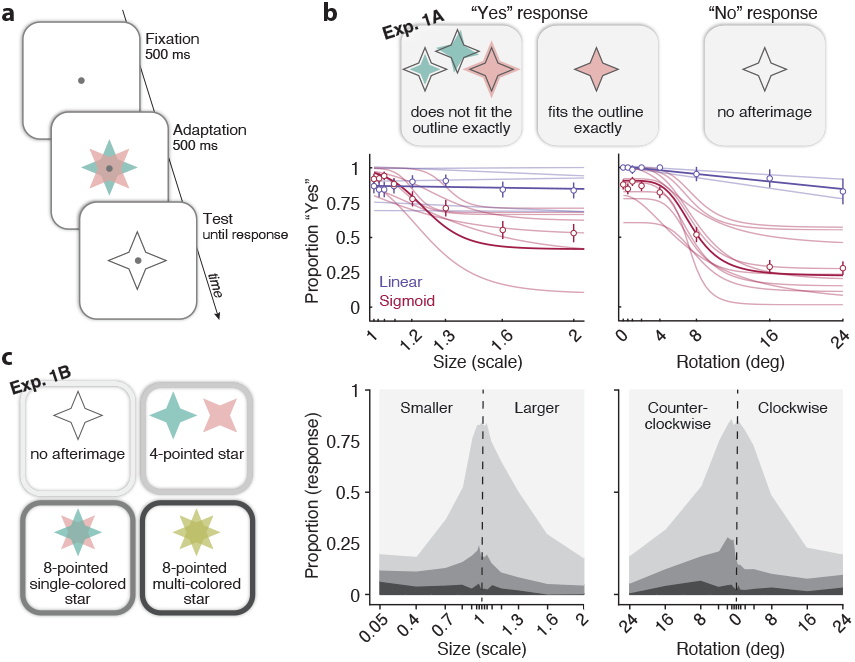
Psychophysical experimental procedure, response options and results for Exp. 1A and 1B. (a) Stimulus and general procedure for Exp. 1A and 1B (top-left). Each trial began with a white screen and a fixation dot. After a 500 ms fixation window, the adapting stimulus was presented for 1 sec. Subsequently, the test outline was displayed until the participant responded. (b) Response options (top-right) and results (below responses) for Exp. 1A. Results are represented as the proportion of responses indicating the presence of an afterimage as a function of the size (left) or rotation (right) of the test outline relative to the size and rotation of the adapting stimulus. Participant responses were fit with either a sigmoid (red fit, n=7 or n=10 for size and rotation, respectively) or linear function (blue fit, n=5 or n=2 for size and rotation, respectively). Participant data is represented by less saturated fits, whereas aggregate data (including SEM) is represented by fully saturated fits. (c) Response options (bottom-left) and results (bottom-right) for Exp. 1B. Results are represented as the proportion of each response as a function of the size (left) or rotation (right) of the test outline relative to the size and rotation of the adapting stimulus (vertical dashed line).

Seven out of twelve participants in the size condition and ten out of twelve participants in the rotation condition exhibited characteristic psychophysical responses (Fig. 3b), which we fit with a sigmoidal function. These participants consistently reported perceiving an afterimage when the deviation in size or rotation was close to zero, i.e., highly similar to the adapting stimulus (for 75% of the time, size of 1.4 ± 0.1 (SEM) and rotation of 5.6*deg* ± 0.8 (SEM)) and tapered off the farther it deviated. The rest of the participants, however, reported perceiving the afterimage near constantly for size and rotation. We fit these individual data using a linear function (slope = -0.02x; y-intercept = 0.89; average proportion = 0.86) and rotation (slope = -0.007x; y-intercept = 1.01; average proportion = 0.93). Given the variability in participants’ responses, which followed either a sigmoid or a linear function, we included this as a parameter in the model.

### The Model Reconciles With Linear Responders And Makes Novel Predictions

In order to reconcile with the variability in response style (sigmoid vs linear) from Experiment 1A, we tuned the model parameters to qualitatively fit individual participants. Fig. 2b (last two rows) shows model responses for participants with a linear profile (Fig. 3b (blue lines)). This pertains to perceiving the (negative) colors from the adapting stimulus as-is, without filling-in (at overlapping gray area of the initial stimulus), or a filled-in afterimage that corresponds to the interpolation of the two colors from the adapting stimulus. The two different outcomes, arising from two different parameter settings, is equivalent to two sets of people each having slightly different dynamics in shape and color processing, relative to the usual responders with a sigmoid profile.

These were unanticipated, novel predictions from the model. However, remarkably, some participants in Experiment 1A reported perceiving such afterimages during debriefing. In Experiment 1B, we validated these predictions.

### Humans Actually See Different Types of Aftereffects

Experiment 1B was designed specifically to test the novel predictions produced by the model. The results of Experiment 1B were similar to those of Experiment 1A: participants reported the presence of an afterimage with less frequency as the size or rotation of the outline deviated from the adapting stimulus (Fig. 3c). To determine whether participants perceived the predictions made by the model, we focused on the data where we expected the greatest proportion of afterimage responses (size = 1; rotation = 0). At this size and rotation, we report with 95% confidence that our participants perceived the 4-pointed star 63.7±2.6% of the time. Additionally, the 8-pointed multi-colored star, was reported 18.2±2.1%, while the 8-pointed single-colored star was reported only 3.5±1.0% of the time. Refer to SI Fig. S3 for more details. Thus, our psychophysical results validate the novel predictions made by our model, suggesting that people perceive the (negative) colors from the adapting stimulus as-is without filling-in (8-pointed multi-colored star) and the interpolation of the two colors (8-pointed single-colored star), indicating slightly different dynamics in shape and color processing across participants.

### Illusion 2: Silencing by Motion

In our second investigation, we examined a visual illusion that reveals an interplay between motion and luminance perception. By modeling and gaining insights for a fairly different phenomenon, we show that AICON has the potential to model broader aspects of human vision.

“Motion silencing” [2], which we call here “silencing by motion” is another striking visual illusion in which objects dynamically changing in luminance (or other features) appear to change slowly or entirely stop changing when they move (refer to SI Movie S5). This silencing-by-motion effect has been shown to scale with speed [2, 14]: The faster a group of dots rotate around a fixation target, the slower the perceived changes in those dots. While various types of changes have been investigated, we focus on modulations of flicker perception—that is, the silencing of changes in luminance by motion.

### Silencing From The Model’s Perspective

A computational model was created to capture the effect of silencing by addressing the interaction between motion and luminance perception. The model, shown in Fig. 4a, estimates the motion and luminance for each object (here dots) in the stimulus, provided as a sequence of images. The output “percept” of the model represents the current estimate of luminance for each dot, which can also be visualized as a sequence of images.

**Figure 4:**
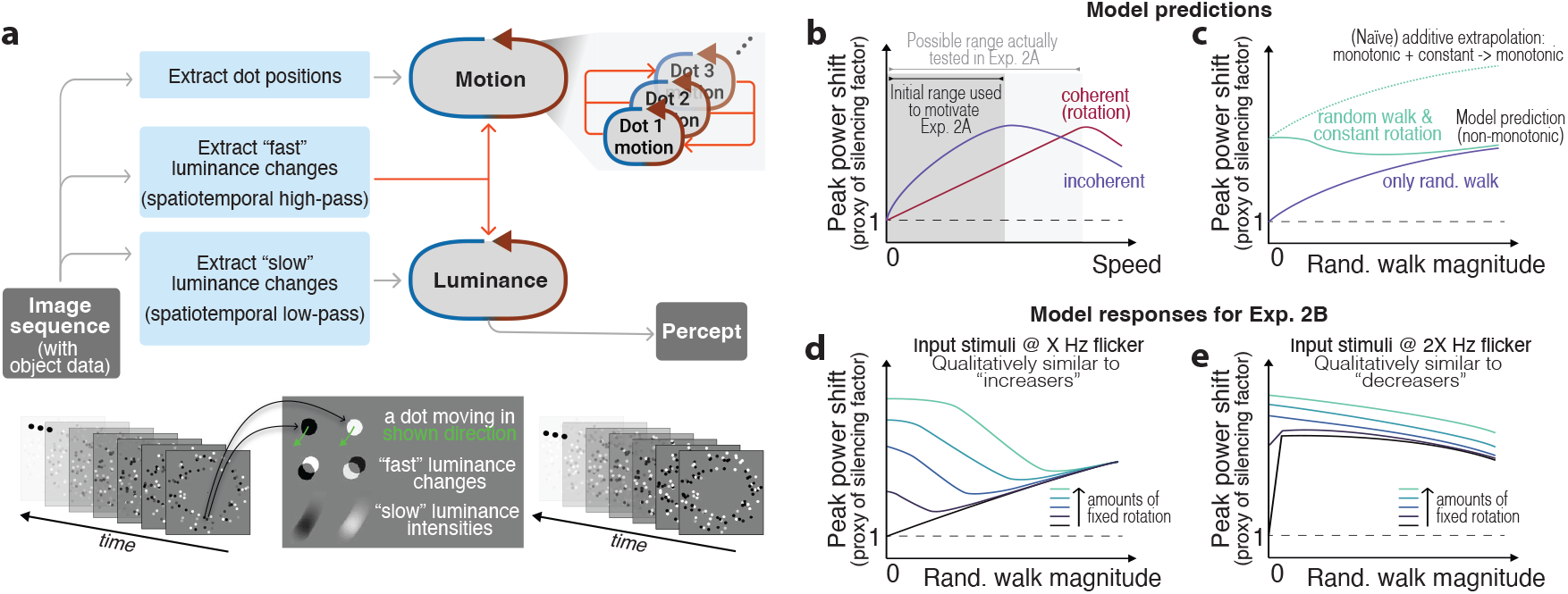
The model for “silencing by motion” reproduces the illusion as seen in humans and makes novel predictions. (a) The model takes a sequence of images as input and uses interactions between recovering motion and luminance quantities to produce the “percept” as seen in humans. The inset gray box illustrates how a moving dot produces motion streaks (“slow” luminance intensities) as well as (“fast”) luminance changes on the image-sensing surface. (b) In the gray interval, the model predicts greater silencing for incoherent motion relative to coherent motion. The light gray interval (and beyond) shows the model’s response for a wider speed range, which was eventually consistent with Exps. 2A and 2B. Silencing in the model is measured through the peak power shift of the spectrum of dot luminance intensities in the “percept” relative to the input. (Find more details in SI Fig. S2.) (c) Model responses show a counter-intuitive non-monotonic trend for combinations of coherent and random walk (incoherent) motion. (d, e) The model qualitatively reproduced the trends observed in Exp. 2B for the two groups (“increasers” and “decreasers”), by varying the input stimulus frequency, which translates to subjects with different internal time constants.

To understand one of the key mechanisms behind silencing, let us first consider the luminance estimation part of the model. Perceptual systems recovering the luminance of moving objects (in this case, dots) encounter a fundamental constraint: A moving dot produces a motion streak on the image-sensing surface [15–18], similar to Fig. 4a (gray box). Consequently, directly-sensed luminance becomes unreliable when motion is present. To overcome this constraint, the model relies on another measure that is contingent on the same motion signal: luminance *changes*. A fast-moving dot will produce a greater luminance transient at a given image-sensing location compared to a slow-moving dot of the same luminance. In essence, the movement of a dot produces a luminance change directly consistent with the amount of motion. Thus, the model uses information from these two sources: “slow” directly-sensed luminance and “fast” luminance changes. The degree of silencing, or the effective change in luminance of a dot, is influenced by the relative contribution of each of these sources and the interactions between motion estimation.

The model estimates the motion of each dot from correspondences over time. In addition to tracking individual dots, the model takes inspiration from the Gestalt principle of common fate [19, 20] to enable motion estimation of the whole (ring of dots, or “donut”). When dots in a neighborhood exhibit similar motion, their individual confidence in motion estimation may be augmented, and through many such mutual interactions, the motion perception of the whole is enhanced.

As illustrated in Fig. 4a, motion and luminance estimation processes for each dot in the stimuli is expressed through Recursive Estimators, and all interactions between processes are encoded with Active Interconnections. (For further details on model, see SI Sec. S2.)

In the beginning of our investigation, this model produced a response similar to previous reports: a monotonous increase in silencing with increasing rotation speeds [2, 14]. Following this, we were able to make a prediction from the model for a type of motion that was, to our knowledge, not tested before in humans. As a consequence of allowing motion estimation of dots to interact with each other, we found that the model produced greater silencing at lower relative speeds when individual dots did not move coherently (see Fig. 4b, gray interval). To determine if this phenomenon holds true in human vision, we psychophysically measured the amount of silencing for coherent and incoherent motion.

### Humans Experience Greater Silencing for Incoherent Motion

In Experiment 2A, twenty participants viewed 100 dots (radius 0.35 dva), randomly distributed in a “donut” shape at an eccentricity of 5 dva to 8 dva around fixation (Fig. 5a), flickering at 1 Hz. After a duration of 3 s, the dots started moving at speeds ranging from 3.75 deg*/*s to 120 deg*/*s—while still flickering at 1 Hz—either coherently around fixation or incoherently in different directions (but not leaving the outline of the “donut”). (Refer to SI Movie S6 for an example of incoherent motion.) Following an additional 3 s, the dots stopped moving, and participants adjusted the flicker frequency using arrow key presses to match the perceived flicker frequency during the movement phase. Participants were allowed to review the movement phase at their leisure by using a key-press to evaluate their adjustment.

**Figure 5:**
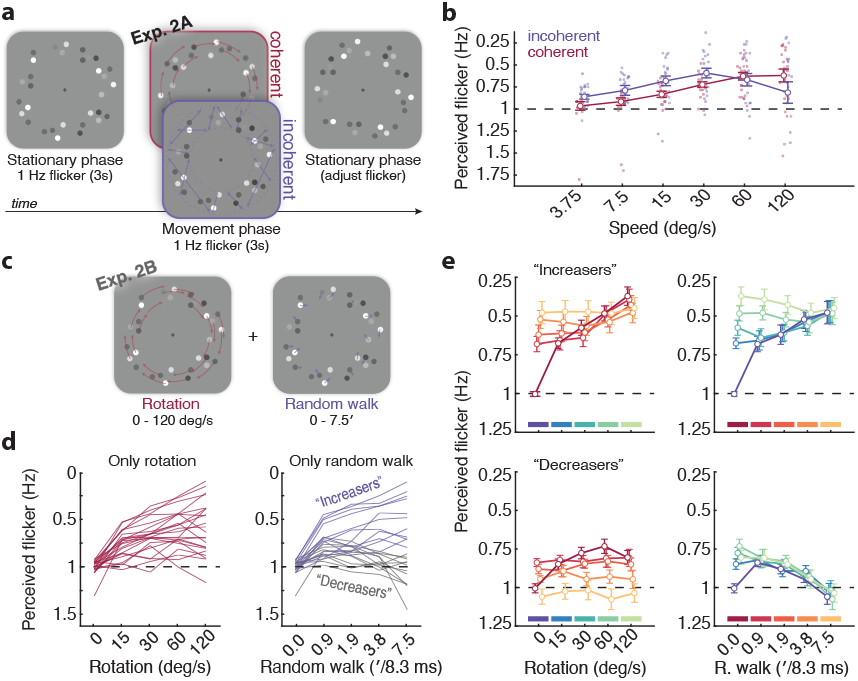
Psychophsysical experimental procedure and results for Exp. 2A and 2B. (a) Stimulus and experimental procedure for Exp. 2A. Dots, arranged in a “donut” shape around the central fixation dot, flickered at 1Hz. After a 3s stationary phase, the dots started moving coherently or incoherently for 3s. In the following stationary phase, participants adjusted the flicker to match the perceived flicker during the movement phase. (b) Group-averaged perceived flicker as a function of speed for coherent and incoherent motion in Exp. 2A. Dots depict individual participants’ averaged measurement, error bars show the standard error of the mean (SEM). Note that the y-axis is inverted, to reflect increased silencing with decreased perceived flicker. Incoherent motion elicited stronger silencing at lower speeds, but this effect reversed at higher speeds. (c) Stimuli in Exp. 2B, combining coherent rotation and random walk motion. (d) Individual participants perceived flicker as a function of pure rotation speed (left) and pure random walk amount (right). The latter was used to divide participants into “increasers” and “decreasers”. (e) Group-averaged perceived flicker for “increasers” (top) and “decreasers” (bottom), showing distinct patterns as a function of rotation speed (left; with different amounts of random walk) or random walk amount (right; with different rotations speeds). Error bars show the SEM. The model successfully captured the non-monotonic trend observed for “increasers” (top-right). Upon changes in model input parameters, the qualitative trends in “decreasers” (bottom-right) were also captured.

In line with previous work [2], participants perceived a lower flicker frequency with higher coherent motion speed (note that the y-axis in Fig. 5(b,d,e) is inverted), indicating that silencing increased with speed. A repeated measures ANOVA with the factors motion type (coherent vs. incoherent) and motion speed (3.75, 7.5, 15, 30, 60 and 120) deg*/*s showed a significant main effect of motion speed [F(1,19) = 1.55, *p* = 0.020]. The main effect of motion type was not significant [F(5,95) = 5.22, *p* = 0.231]. However, we observed a significant interaction effect of motion coherence as a function of motion speed [F(5,9) = 9.10, *p <* 0.001]. In line with the predictions of our model, at relatively lower speeds, incoherent motion elicited greater silencing than coherent rotation. This effect reverses at higher rotation speeds (60 deg*/*s, Fig. 5b), which was not initially predicted by the model, but could later be reconciled.

### The Model Makes Novel Predictions for Combinations of Coherent and Random Walk Motion

Independently, without making any changes to the model, we obtained further predictions for untested conditions. Preliminary data from Experiment 2A suggested that the model predictions were indeed accurate—showing greater silencing for incoherent motion relative to coherent motion. Consequently, we sought to investigate how silencing would be influenced when both coherent and incoherent motion are present at varying magnitudes (see SI Movie S7– S8 for example stimuli). Note, the type of incoherent motion we tried here is different from that in Experiment 2A. Instead of each dot moving in a straight path of a randomly chosen direction, it undergoes a “random walk” to a specified distance per frame. Such motion allows us to create combinations of coherent and incoherent motion. In the following, we delineate this *type* of incoherent motion as “random walk motion”.

When the model was exposed to stimuli with different magnitudes of random walk motion added to some constant magnitude of coherent motion, we observed a non-monotonic trend in the model’s response (Fig. 4c). That is, for some combination of speeds of coherent and random walk motion, the degree of silencing could be *lower* than purely coherent or purely random walk motion. This result was initially counter-intuitive, as the naive addition of two monotonic trends would result in another monotonic trend. To validate this unexpected prediction, we designed Experiment 2B.

### Humans Show Idiosyncratic Differences For Various Types of Motion

In Experiment 2B, we systematically manipulated the amount of motion coherence by adding different magnitudes of displacement (0-7.5 arcmin/8.3 ms; i.e., per refresh @120 Hz) to different rotation speeds (0-120 deg/s) (Fig. 5c). As in Experiment 2A, participants (N = 23) adjusted the perceived flicker speed during a stationary phase to match the 1 Hz flicker during the movement phase.

When inspecting the data, it became obvious that the effect of motion on perceived flicker varied profoundly among participants (Fig.5d). On the group level, silencing increased with both rotation speed as well as random walk magnitude. However, at the individual observer level, for some participants this effect was strongly pronounced, whereas for others it was even reversed. We fit a linear regression model to each participant’s data and tested the distribution of the obtained slopes using a Shapiro-Wilk test, which indicated that the distribution of slopes in the “Pure random walk” condition (Fig. 5d, right) departed significantly from normality [W = 0.91, *p* = 0.0346]. To explore the opposing trends, we split our sample into two groups: “increasers” (n = 11) with slopes above the group median of the “Pure random walk” condition (Median = -0.016), and “decreasers” (n = 12) with slopes below.

For the “increasers”, silencing monotonically increased with rotation speed when no random walk was added (Fig. 5e; top left, red curve), in line with Experiment 2A and previous work [2]. A similar trend was observed for increasing random walk magnitudes in the absence of rotational motion (Fig. 5e; top right, blue curve). Interestingly, despite both motion types individually displaying a monotonically increasing trend in silencing, their combination resulted in a non-monotonic pattern (Fig. 5e; top right, light green), as predicted by the model (Fig. 4d).

In comparison to previous research [2], the “decreasers” showed an unexpected trend in silencing: the amount of silencing decreased with increasing random walk motion in the absence of rotation motion (Fig. 5e; bottom right, blue line) as well as for any amount of added rotation motion. Interestingly, this inverse relation can also be observed in pure rotation trials without random walk (Fig. 5e; bottom left, red curve): Instead of the typically reported monotonic increase of motion silencing with speed (also observed for “increasers” and in Experiment 2A), motion silencing only increases up to a rotation speed of 60 deg/s. In fact, a similar trend was observed in Experiment 2A, where motion silencing either plateaued (for coherent motion) or even declined (for independent motion) at the highest speeds.

### The Model Reconciles With Unexpected Responses in Humans

In an attempt to obtain clarity into the varied trends observed in the two groups, we subjected the model to various conditions. Surprisingly, we found that both groups can be captured using only a single parameter change. Specifically, the model produced qualitative trends that aligned with both groups for two different input stimulus frequencies, akin to models instantiated with two distinct “time constants”. Moreover, at intermediate frequencies, the model produced trends that exhibited a continuous transition from one group to the other. These finding suggests that participants with different internal dynamics perceive the illusion in distinct ways.

## Discussion

We demonstrated that models using AICON generated novel and experimentally valid predictions, and even guided the design of further psychophysical experiments. In both the phenomena we investigated, the models also helped explain variance between subjects. Even though AICON was originally developed for robotic applications, our results highlight its promising potential to model and gain insight into various aspects of human vision. This concept may seem too aspiring at first, but we argue that AICON possesses unique qualities that make it particularly well-suited for studying biological information processing. In the following, we elaborate on these arguments and discuss the broader implications and limitations.

A common approach in computational modeling is to “abstract away” the substrate-level details to leverage the stronger capabilities afforded by abstract representations. In this regard, AICON provides a “mid-level” abstraction that is not bounded by physiological constraints. However, unlike many approaches, this abstraction is not merely a theoretical construct existing in a void, nor is it just another modeling framework chosen solely for its mathematical elegance or compatibility with a particular computing infrastructure.

A key tenet underlying AICON’s abstraction is the recognition that rich interactions and interdependencies between perceptual/motor processes are likely a hallmark of intelligent information processing [3, 12]. Biological intelligences did not evolve as neatly decomposable modular systems, nor did they completely discard modularity. Instead, biological intelligences can be viewed as “weakly decomposable” systems with some degree of modularity, where the modules maintain rich interactions with each other. AICON encodes this crucial principle of “weak decomposability”, and manifests as the ability to break down a complex problem into semi-isolated subproblems (Recursive Estimators), while still allowing their solutions to mutually inform and constrain (Active Interconnections) each other over time.

Furthermore, this abstraction is grounded in robotics, where the primary goal is to generate robust artificial behaviors for real-world interaction. Viewed from this perspective, the abstraction inherits key principles shaped by the same constraints that biological agents must overcome: efficiently extracting requisite information from noisy sensory and motor variables, by possibly leveraging several sensory and sensorimotor regularities, and interactions between those regularities. Consequently, this abstraction incorporates features shaped by the demands of generating robust behavior in real-world contexts. Such demands were possibly an important evolutionary pressure that molded biological information processing.

Abstractions like AICON have likely remained unexplored in the context of biological information processing due to three main reasons: 1) existing abstractions do not sufficiently emphasize the crucial role played by interactions between different modules; 2) existing abstractions lack a focus on design principles that ultimately facilitate robust real-world behavior; and 3) using an unconventional abstraction (such as AICON), at least in the beginning, requires a concerted interdisciplinary effort [21].

Having discussed the principles that underlie AICON, let us now take a look at how AICON is valuable to study human vision.

Recall that AICON consists of two functional elements: Recursive Estimators and Active Interconnections. Recursive Estimators model dynamics and perform inference over the Active Interconnections. In this work, we only resort to Bayesian inference, but other schemes may be used as we explain later. Nevertheless, modeling dynamics through recursion and performing (Bayesian) inference is a powerful signal processing scheme: It helps in rejecting noise, recovering required information from minimal data, and adapts quickly when conditions change drastically [13]. As a collective of multiple Recursive Estimators and Active Interconnections, they leverage the same properties in a combined but highly complex space.

The main reason behind this ability is that AICON places Active Interconnections—the interactions between parts of a system—in a central role. In the models we presented here, the interactions between multiple rather straight-forward perceptual processes (e.g., color processing, motion estimation, etc.) manifested as complex but non-trivial behavior. This was also validated in humans phenomenologically, which implies that such interactions between basic perceptual processes may play a major role in human vision [4, 22].

Further, AICON offers an explicit balance between modularity and interconnectedness. Modularity (through Recursive Estimators) is helpful to encode some well-known properties of a (sub-)system, as well as for reusability. The gaps left by parts of a system that are hard to fully specify are filled by the Active Interconnections between the modules, as they express some mutual information that are typically less restrictive to specify. This balance offered by AICON allows us to quickly create a model and incrementally adjust it to fit to the needs.

For the future, AICON’s grounding in efficient sensorimotor integration for behavior will facilitate the modeling of more complex aspects of human information processing. That means that with AICON, our research is not limited to studying visual inferences in illusions alone. We can also explore processes such as (eye-movement-mediated) attention, adaptation, cue integration etc., in more ecologically relevant settings that involve real-world interactions and human behavior.

However, every solution to modeling faces a fundamental trade-off. Since AICON is a “mid-level” abstraction, it offers the ability to rapidly create models with a large scope. This advantage comes at a cost of reduced resolution in capturing biological mechanisms—at least in the early stages of study. This contrasts with another common approach to studying biological information processing: bottom-up modeling. Bottom-up approaches often provide better physiological fidelity but tend to capture phenomena with a narrower scope. Let us examine this trade-off in more detail for each case.

In our work, the abstraction manifests as a need to resort to qualitative comparisons when comparing the models’ performance against human data. This primarily stems from the disconnect to human physiology. For example, the sense of time in the models arises solely from the update rate of each Recursive Estimator, which is not grounded in the actual temporal dynamics of the modeled process. As a result, measures involving time such as decay time and frequency, are arbitrarily scaled in the models. The same applies to various other aspects, such as the choice of color space, scale of luminance, scaling of spatial coordinates. Nevertheless, we can rely on qualitative predictions and trends. For example, if increase in value of a stimuli parameter elicits a weaker response in the model, the same trend should hold qualitatively in humans. In both the phenomena we investigated here, the models’ qualitative predictions and trends captured essential aspects of the studied phenomena.

In contrast, bottom-up approaches provide more realistic predictions often at the cost of scope of the modeled mechanism. This can lead to mutually contradictory accounts of the same mechanism. For example, in the sub-area of early visual motion perception (areas V1 and MT), two notable models that extend [23] or draw inspiration [24] from the seminal spatiotemporal energy model [25] propose fundamentally incompatible sub-mechanisms. The former [23] advocates for pooling responses of V1 (simple) cells, while the latter [24] suggests directly integrating from two types of V1 cells. Such instances of competing theories, often unresolved, can be found in many bottom-up modeling approaches. This maybe a consequence of trying to maintain physiological/neuroanatomical accuracy despite the brain’s incredible complexity. This, in turn, biases researchers to resort to modeling assumptions that risk overspecifying aspects that are not fully understood.

We believe that, in the early stages of studying biological information processing, it is advantageous to accept the trade-off an abstraction like AICON offers. This approach allows for greater flexibility in changing directions when certain observations do not hold true or in reconciling seemingly incompatible explanations. By picking an appropriate abstraction, fewer pivots, reconciliations, or overspecifications may be necessary in the long run. And as we gather a substantial number of working mechanisms, we can gradually anchor some processes to physiological constraints, making comparisons to human data more concrete and quantitative. This iterative process would eventually lead to a deeper understanding of biological information processing while maintaining a balance between scope and fidelity.

It also behooves us to compare our approach with another “mid-level” abstraction: Bayesian computational models. These are popular in human vision and they model large parts of vision under Bayesian inference schemes [26–32]. Like our approach, Bayesian computational models avoid a bottom-up approach and resort to a “mid-level” abstraction. Additionally, the models we presented in this paper and Bayesian computational models employ Bayesian inference in some form. Despite these similarities, there are key differences between our approach and typical Bayesian computational models.

While we currently use (modified) Bayes filters as Recursive Estimators that in effect perform Bayesian inference over Active Interconnections, AICON is not restricted to Bayesian inference. Recursive Estimators can be seen as a class of estimation techniques that leverage their recursive property to impose priors on recovering quantities of interest. Although Bayes filters are a particularly elegant instance of this [13], we could also use other variants, such as Recursive Least Squares (RLS), Least Mean Squares (LMS), and Auto-Regressive Moving-Average (ARMA) [33–35].

Many works also commit to hierarchical architecture [30, 32], but AICON applies to arbitrary topologies. A hierarchical topology can be a strong inductive bias. After all, the broad organization of the visual cortex also exhibits a hierarchical structure [36, 37], and some approaches show hierarchy leads to useful functions such as “learning” concepts of abstraction [29]. While we do employ hierarchy when needed, such as in the robotic perception system [3] in Fig. 1, it is not a central focus of our approach. Instead, we emphasize the importance of interactions between multiple processes, which may manifest as hierarchical, cross-modal or other types of relationships.

In the following, we briefly discuss limitations that arise from simplifications in the presented models as well as in modeling capabilities of AICON.

The model for “Illusion 1: Fill-in color after-effect illusion” produces positive-colored afterimages instead of negative-color afterimages experienced by humans. To elicit negative-colors, we could have optionally added a mechanism (not a Recursive Estimator) that scales inputs with a pixel-wise dynamic gain. However, we left this out as such mechanisms are secondary to understanding the phenomenon of interest: interaction between color and shape perception.

The motion estimation process in the model for “Illusion 2: Silencing by motion” relies more heavily on correspondences from object positions over time—second-order motion [38], than on motion information obtained through its interconnection to luminance estimation—a type of first-order motion perception [38]. In human vision, these two types of motion perception likely interact in more complex ways, possibly with inputs from other first-order motion cues similar to spatiotemporal motion energy [25].

In both models, we specify all their aspects without “learning” from data. While parts of the models could potentially be trained using data [3], we consider it more prudent at this nascent stage of studying biological information processing to first assemble a few candidate “canonical” mechanisms that are some combinations of Recursive Estimators and Active Interconnections. Once a few such mechanisms are available, data-driven approaches in this constrained context could prove to be more efficient and stable.

With the two functional elements of Recursive Estimators and Active Interconnections, we presently lack a clear path to implement high-level cognitive functions. However, certain high-level functions may be achieved as an emergent property from the functionality offered by AICON. Some studies of visual working memory [39–41] suggest that sensory cortical areas play a crucial role in maintaining working memory representations. Processes undertaken by sensory cortical areas seem to be amenable to AICON, as we demonstrated in the models here. For other cognitive functions, we may also need to create other functional elements in the future. Nevertheless, as demonstrated in this paper, certain aspects of human vision can currently be modeled solely through the existing feature-set of AICON. Since several perceptual phenomena exhibit interactions between basic perceptual processes, we believe that, for now, many of these can be directly analyzed using AICON.

Having discussed limitations of our approach, let us now make some concluding remarks highlighting the potential of a useful abstraction suitable for studying biological information processing.

It may seem surprising that the models we presented here capture many important aspects of the studied phenomena. However, as we advocated earlier, we think this capability of the models arises from using an appropriate level of abstraction to study perceptual problems. This insight can be valuable for many problems beyond the scope of our study here and imply an alternative systematic way to “decipher” the mechanisms underlying human information processing.

Using AICON, or abstractions similar to AICON, we can quickly create a number of different mechanisms representing different aspects of human perception. But this abstraction extends to establishing interactions *between* those mechanisms. When setting up interactions, some may be easier to characterize and some interactions may be more subtle and involve experimentation. And some interactions may lead to beneficial higher-order couplings. Using AICON we encountered all these kinds of interactions within the models we presented, and also found it suitable to experiment with systematically.

Thus one can use an abstraction to build larger and larger models of the perceptual system, starting with smaller mechanisms, followed by compositions of several such mechanisms, and so on. In the context of robotics, AICON has been used to build such compositions [42, 43]. As we mentioned earlier, as a consequence of the abstraction, at first we might only be able to rely on qualitative comparisons with human performance, but as our understanding grows larger, we can anchor some processes to physiological constraints and slowly make the comparisons more concrete.

Having made the case for exploring human perception at a useful level of abstraction, we must emphasize that we are not advocating for stopping studies at other levels of abstraction. Insights obtained through, e.g., studies done at the physiological level can inform certain constraints a model at the level of abstraction we study ought to follow, and insights obtained from such models can guide studies at a different abstraction. While we (as a community) span the study of intelligent information processing in various abstractions, we would like to highlight that large advances can be made in an interdisciplinary study when we have a common abstraction “language” similar to our work.

## Methods

### General Psychophysical Setup

All human experiments were executed using a PC with an Ubuntu 20.04 operating system. Stimuli were projected using a PROPixx (VPixx Technologies, Saint-Bruno, QC, Canada) with a 120 Hz vertical refresh rate and a resolution of 1920 x 1080 pixels. Stimuli were presented on a 16:9 (150 x 84 cm) video-projection screen (Stewart Luxus series ‘GrayHawk G4’, Stewart Filmscreen, Torrance, CA), mounted on a wall, 180 cm in front of the participant. The experimental code was written in MATLAB (Mathworks, Natick, MA, USA) with Psychophysics and Eyelink toolboxes [44, 45]. Fixation was checked online using the Eyelink 1000+ tabletop system (SR Research, Osgoode, ON, Canada) at a sampling rate of 1000 Hz.

Participants were recruited through university mailing lists and general lines of communication (e.g., social media and word of mouth). Exclusion criteria for our studies included: (1) corrected visual acuity worse than 20/20 or (2) any form of color deficiency (i.e., red-green or blue-yellow partial color blindness) or (3) neurological conditions. The protocols were approved by Humboldt-Universität zu Berlin’s Institutional Review Board and our methods adhered to the Declaration of Helsinki (2008). Experiment 1A and 1B (https://osf.io/ap8yk) as well as Experiment 2B (https://osf.io/g453q) were preregistered.

### Experiment 1A

Twelve adults (females: 5, age range: 20-30) with normal or corrected-to-normal visual acuity and no color vision deficiency, participated in the study after providing written informed consent.

### Stimuli

The adapting stimulus consisted of an eight-pointed colored star comprised of two four-pointed stars (one reddish, RGB: [226 163 163] and the other cyan, RGB: [116 190 183]) with a gray center covering the intersection of the two four-pointed stars, modeled after the adapting stimulus in Van Lier et al. [1] (Fig. 3(a)). The adapting stimuli had a visual angle of (7.21 dva) and was presented at full color. The test outline was presented at various sizes (1, 1.0274, 1.0488, 1.0954, 1.1832, 1.3416, 1.6125 or 1.9494 times the area of the adapting stimulus) and orientations (0, 0.5, 1, 2, 4, 8, 16 or 24°) with respect to the adapting stimulus. Both the adapting stimulus and test outline were presented against a white background with a small fixation dot (0.19 dva) at the center. The adapting stimulus was always presented in the same size (7.21 dva) with any possible positional jitter combination in the range of [-1.19 to +1.19] dva from the screen center (in x and y) and with an initial randomized orientation [0 to 30°] clockwise or counterclockwise from center across all trials. The test outline was presented with: (1) a size change, or (2) a change in rotation, randomly interleaved either clockwise or counterclockwise.

### Procedure

Each session (total two) consisted of two randomized blocks: (1) size change only and (2) rotation change only. On each trial, participants fixated the dot for 500 ms before the adapting stimulus was presented for 1 sec. Subsequently, the test outline appeared and remained on the screen until participants reported what they saw using one of three arrow keys: the afterimage did not fit the shape exactly (left arrow), there was no color afterimage (down arrow), or the color afterimage fit the shape exactly (right arrow).

### Data Analysis

Each participant produced a total of 512 trials, made up of: two conditions (size, rotation), two sessions, eight stimulus levels, and 16 trials. To determine whether the outline elicited an afterimage, we binned responses for “the color afterimage did not fit the shape exactly” and “the color image filled the shape exactly” as a “yes” and the response “there was no color afterimage” as a “no”. Our attempt to fit a psychometric curve to response data for each participant and condition across two sessions revealed that participant responses could be categorized in two groups: (1) participants who responded linearly (decreasing their response to seeing an afterimage as the shape outline deviated from the adapting stimulus) and (2) participants who responded closer to a step function (reporting the presence of an afterimage up until a certain deviation and then reporting no afterimage). For participants who were fit with psychometric curves, we used the Palamedes Toolbox [46].

### Experiment 1B

Twenty adults (females: 14, age range: 19-32) with normal or corrected-to-normal visual acuity and no color vision deficiency, participated in the study after providing written informed consent.

### Stimuli

The adapting stimulus for Experiment 1B was the same as for Experiment 1A (Fig. 3(a)). However, there were differences in the size of the test outline. In addition to the test stimulus being presented at a size change of 1 to 2 times the area of the adapting stimulus, we also presented the test outline from 0 to 1 times the area of the adapting stimulus for a total of 15 sizes. The change in rotation remained the same as in Experiment 1A.

### Procedure

The procedure for Experiment 1B was similar to Experiment 1A, with only a few key differences. First, each session was completed in two blocks of randomized, interleaved conditions of (1) size change and (2) rotation change. Second, the response options were different. Following the test outline, participants reported their immediate percept using one of four arrow keys: there was no afterimage (down arrow), the afterimage was a 4-pointed star (up arrow), the afterimage was a multi-colored 8-pointed star (right arrow) or the afterimage was a single-colored 8-pointed star (left arrow).

### Data Analysis

Each participant produced a total of 736 trials, made up of: 23 stimulus levels (15 for size and eight for rotation), two sessions, and 16 trials. To validate model predictions and determine if participants indeed perceive the different after-images, for each response, we computed the 95% binomial confidence intervals at the stimulus where we expected the greatest proportion of “yes after-image present” responses. Meaning, where the size and rotation of the outline is equal to that of the adapting stimulus) (total trials = 1280).

### Experiment 2A

Twenty adults (females: 8, age range: 20-63) with normal or corrected-to-normal visual acuity and without color vision deficiency, participated in the study after providing written informed consent.

### Stimuli

The stimuli were modeled from Suchow and Alvarez [2] and consisted of 100 dots randomly distributed in a “donut” shape – a ring with a 5 dva inner radius and a 8 dva outer radius. Each dot (radius 0.35dva) was randomly assigned a starting gray value with a corresponding luminance between 48 and 118 cd*/*m^2^, which flickered sinusoidally at 1 Hz, with an amplitude of 26 cd*/*m^2^. The stimulus was presented against a gray background with a small fixation dot (0.187 dva) at the center. The position of the stimulus was randomly jittered in the range of [-1.19 to +1.19] dva from the screen center.

### Procedure

Each trial consisted of three phases: (1) stationary phase, (2) movement phase, and (3) the matching phase (Fig. 5a). In the stationary phase, the stimulus was presented without movement for three seconds. It was then followed by another three seconds with rotational or translational movement at velocities of 3.75, 7.5, 15, 30, 60 and 120 deg per second. During both the stationary and the movement phase, dots flickered sinusoidally in luminance at a rate of 1 Hz. Lastly, in the matching phase, participants were asked to adjust the flicker frequency of the stationary phase to match the perceived flicker frequency of the movement phase, which also had an initial flicker frequency of 1 Hz. Participants were allowed to switch between the stationary and the movement phase while adjusting the flicker frequency until they were satisfied with the match. We varied movement velocities between trials, and movement type (rotational or translational) between blocks.

### Data Analysis

Each participant produced three trials per speed and condition. Thus for each speed and condition, we averaged across a total of 60 trials, made up of twenty participants and three trials. Fig. 5b shows the group average of these measurements per motion type, error bars depict the standard error of the mean (SEM). To assess statistical significance, we used repeated measures ANOVAs. If the sphericity assumption was not met, we report Greenhouse-Geisser corrected *p*-values.

### Experiment 2B

Twenty three adults (females: 11, age range: 19-39) with normal or corrected-to-normal visual acuity and no color vision deficiency, participated in the study after providing written informed consent.

### Stimuli & Procedure

Dots in Experiment 2B had a size of 0.7 dva; number, arrangement, and flicker properties were identical to Experiment 2A. The trial sequence differed as follows: Each trial consisted of two phases, starting with a 3s movement phase (flicker frequency: 1 Hz), followed by a stationary phase (initial flicker frequency randomly selected between 0.001 and 2 Hz), during which participants adjusted the flicker to match the perceived flicker frequency during the preceding movement phase. They now used a turning knob for their adjustment. As in Experiment 2A, participants were allowed to switch between the stationary and the movement phase while adjusting the flicker frequency until they were satisfied with the match. Dot motion during the movement phase was composed of different rotation speeds (0, 15, 30, 60, 120 deg/s) with different amounts of added random walk (0, 0.9, 1.9, 3.8, 7.5 arcmin/8.3 ms), randomly intermixed within each experimental block. Data collection was divided in two sessions per participant, each comprising 4 blocks with 25 trials each (one per rotation speed - random walk amount combination).

### Data Analysis

Each participant produced 200 trials, eight per rotation speed - random walk amount combination. We first averaged individual participants’ measurements per condition. Line plots in Fig. 5e show the group average of these measurements, with error bars depicting the standard error of the mean (SEM). To assess statistical significance, we used repeated measures ANOVAs. If the sphericity assumption was not met, we report Greenhouse-Geisser corrected *p*-values.

## Supporting information

"Illusion 1: Fill-in Color After-Effect Illusion": Demo movie

"Illusion 1: Fill-in Color After-Effect Illusion": Example stimuli used in experiments. Condition: No rotation, no size change

"Illusion 1: Fill-in Color After-Effect Illusion": Example stimuli used in experiments. Condition: 7 deg rotation, no size change

"Illusion 1: Fill-in Color After-Effect Illusion": Example stimuli used in experiments. Condition: No rotation, 120% size change

"Illusion 2: Silencing by Motion": Demo movie

"Illusion 2: Silencing by Motion": Example stimuli used in Exp. 2A (incoherent motion)

"Illusion 2: Silencing by Motion": Example stimuli used in Exp. 2B (only random walk motion)

"Illusion 2: Silencing by Motion": Example stimuli used in Exp. 2B (random walk motion & coherent motion)

## Data Availability

Raw data will be available on the Open Science Framework database (https://osf.io/h6k38/) upon publication.

## Code Availability

The data analysis code will be available under the same Open Science Framework database (https://osf.io/h6k38/) upon publication. The model-related code will be available as a single repository on GitHub upon publication.

## Acknowledgements

We would like to thank Maren Eberle, Frederike Fischer, Carmen Haake, Arne Stein and Marida Zhuba for their assistance with the psychophysical experiments. We appreciate Vito Mengers for comments on early versions of this manuscript. Special thanks to Furkan Davulcu for efforts in independently implementing and validating the computational model for “Illusion 1: Fill-in Color After-Effect Illusion”.

## Funding

We gratefully acknowledge funding provided by the Deutsche Forschungsgemeinschaft (DFG, German Research Foundation) under Germany’s Excellence Strategy – EXC 2002/1 “Science of Intelligence” – project number 390523135.

## Supporting Information

### S1 Functional Elements of AICON

In order to interpret the model details provided in later sections of this supporting information, we first describe the functional elements of AICON in detail. In these descriptions, we will also make clear our definitions and conventions, which deviate from the standard usage in a few ways.

There are two functional elements in AICON: Recursive Estimator (RE) and Active Interconnection (AI). We will gradually build up to these elements by first providing some background on REs, followed by the form of RE we use in AICON. Finally, we provide a description for AI.

#### S1.1 RE in General

REs model physical processes using a quantity. A quantity could be multivariate and characterize the physical process through (an approximately) Markovian assumption.

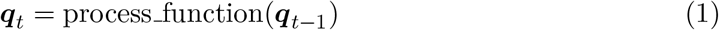

Here, ***q***_*t*_, ***q***_*t*−1_ ∈ ℝ^*n*^ represent the multivariate quantities at two consecutive time steps *t* and *t* − 1 respectively. The function that maps between them, process function(·) : ℝ^*n*^ ↦ ℝ^*n*^ stands for how the “state” of a process evolves in time. One can determine the value of the quantity ***q***_*N*_ at time *t* = *N* by recursively applying process_function(·) to ***q***_*t*_, starting from ***q***_0_(*t* = 0). The choice of parameterization for ***q*** and process_function(·) depends on the process being modeled.

The most important function of an RE is to *infer* ***q***_*t*_ from observations of the modeled process. Observations could also be multivariate and relate to the process as following:

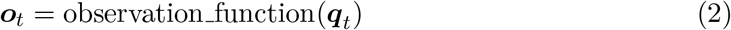

Here, ***o***_*t*_ ∈ ℝ^*m*^ represents an observation of the running process at time step *t*, characterized by observation function(·) : ℝ^*n*^ ↦ ℝ^*m*^.

An example for a quantity ***q***_*t*_ is position and velocity of a moving object in meters at time *t*. The process function(·) could encode a constant velocity heuristic: only allowing jumps in position consistent with its velocity. The process’ observation ***o***_*t*_ could be the position of that object sensed on imager, in pixels. Then the observation function(·) simply encodes conversion from meters to pixels. Over time, an RE can recover the position and velocity of the object from a sequence of observations ***o***_1:*t*_ of the object on the imager.

A Recursive Estimator thus recursively estimates ***q***_*t*_ using observations ***o***, mediated by the functions, process function(·) and observation function(·). Depending on the scheme of Recursive Estimation, only the current observation ***o***_*t*_, a few previous observations ***o***_*t*−*K*:*t*_, or all observations ***o***_1:*t*_ may be used.

#### S1.2 Extended Kalman Filters as REs

There are many recursive estimation techniques. A few examples are Least Mean Square (LMS), Recursive Least Squares (RLS), ARMA and its variants [1, 2]. AICON can include any scheme of recursive estimation. **However, for the rest of the text and the scope of this paper, we only consider a *certain class* of Recursive Estimators: Bayes filters [3]. More particularly, we use a modified form of Extended Kalman Filters that is an instance of Bayes filter**. In this subsection, we will explain the standard form of Extended Kalman Filter, and then detail our modifications in the next subsection.

Please note, in order to conform to conventions used in the literature [3], we perform a change of variables! At time step *t*, the quantity represented by a physical process is called a state, ***x***_*t*_ ∈ ℝ^*n*^ (instead of ***q***_*t*_), and an observation is represented by ***z***_*t*_ ∈ ℝ^*m*^ (instead of ***o***_*t*_).

A Bayes filter provides a structured way to obtain the following probability distribution: *p*(***x***_*t*_|***z***_1:*t*_). That is, the probability distribution of the state ***x***_*t*_ contingent on all past observations, ***z***_1:*t*_. And with Extended Kalman Filters, the probability distribution is Gaussian: *p*(***x***_*t*_|***z***_1:*t*_) = 𝒩 (***µ***_*t*_, **Σ**_*t*_), where 𝒩 (·) signifies a Gaussian distribution with a mean ***µ***_*t*_ and covariance **Σ**_*t*_.

In Extended Kalman Filters, the process function(·) is characterized as:

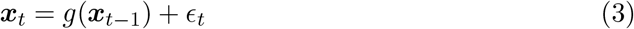

This is called the **process model**, and here a zero-mean Gaussian noise, 𝒩 (**0, *R***), is assumed to disturb the process. A “control input” ***u***_*t*_ can also be included (*g*(***x***_*t*−1_, ***u***_*t*_)), but we drop this for the scope of this work.

And the observation function(·) is given by:

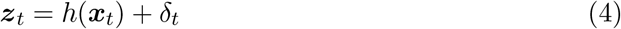

This is called the **observation model**, and a zero-mean Gaussian noise, 𝒩 (**0, *Q***) is assumed to affect the observation of the process that is represented by ***x***_*t*_.

Extended Kalman Filters *infer* ***x***_*t*_ from a series of noisy observations ***z***, mediated by the **process model** and **observation model** and using Bayesian inference.

#### S1.3 REs in AICON

As was noted, for this paper, we restrict to Extended Kalman Filters (EKF) with modifications. In other words, REs in this paper have the specifications as mentioned below.

In our modified EKF, the **process model** retains the same form as the standard form Eqn. 3. For completeness, we restate this:

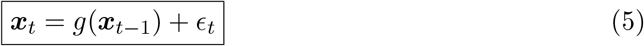

However, we modify the manner in which RE uses observations for inference. In AICON, an RE may be connected to several Active Interconnections (AI), and each AI maybe connected to several REs at once. This manifests as an RE requiring multiple observation models and each observation model taking multiple observations at once respectively. Inference with such inputs cannot be done with the standard EKF as RE, and thus necessitates modifications.

To understand the shift in perspective of inference with AICON, let us first walk through an example and then work up to the general form.

Consider a four RE setup as presented in Fig. S1a. The RE at the center is connected to three other REs using two AIs. (We will detail the form of AI in the next subsection.) REs receive and transmit information through their **ports**, as illustrated in Fig. S1b. For the purposes of this explanation, let us only look at the receiving function of the **ports**.

At **Port 1**, the information coming from the RE on the left serves as an “observation”, say 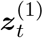 with a covariance (for parameterizing it as a Gaussian distribution) of ***Q***^(1)^. In contrast to the standard EKF (Eqn. 4), we use the following form for the observation model:

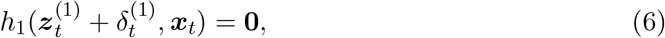

where 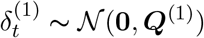. When there is no implicit parameterization between 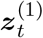 and ***x***_*t*_, this form is equivalent to the standard form (Eqn. 4):

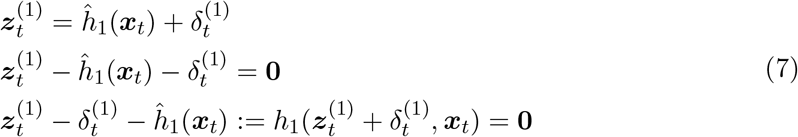

Note, the inversion of sign before 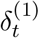 is inconsequential as it is a zero-mean Gaussian.

At **Port 2**, the information coming from the two REs on the right serve as independent “observations”, say 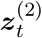 and 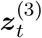, with covariances ***Q***^(2)^ and ***Q***^(3)^ respectively. Following a similar form as before (Eqn. 6), the observation model is given by:

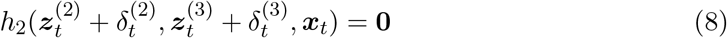

So, the RE illustrated in Fig. S1b has two **ports**. Each **port** is supported by an observation model (Eqn. 6 and Eqn. 8). Both observation models have a non-standard form that allows multiple observations to be consumed by the RE.

Having looked at an example, now we can state the **general form of observation models** for the modified EKF type of RE in AICON. For an RE modeling a physical process using state ***x***_*t*_, exchanging information through *K* **ports**, and each **port** connected to *J* variables, the *k*^*th*^ observation model (*k* = {1, …, *K*}) is given by:

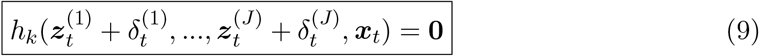

There is an additional feature we incorporate in REs of AICON: they can also optionally reject outliers using a Mahalanobis distance threshold. That is, during inference using *k*^*th*^ observation model, if the amount of adjustment calculated by the inference procedure is higher than a threshold, then all inputs that arrived through the *k*^*th*^ **port** will be discarded.

#### S1.4 Active Interconnections (AI) in AICON

An Active Interconnection (AI) relates multiple quantities being estimated in the REs, and thus serves as a medium for the REs to infer their respective quantities.

Here, we use the same example from Fig. S1 first, before working up to the general form of an AI. Consider **Port 2** of the RE with state ***x***_*t*_ (Fig. S1b). An AI is a function that relates the state of this RE to the states of the connecting REs. The form required by AI can be derived from any closed-form expression, say:

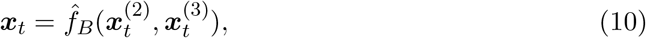

where 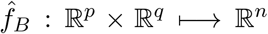, and 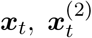 and 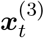 are states being estimated by the corresponding REs as shown in Fig. S1a.

This can be reformed as:

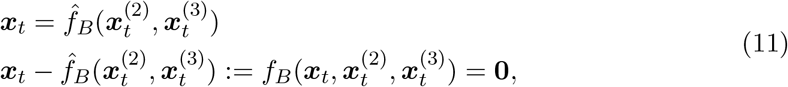

which is the same expression mentioned in Fig. S1a for the corresponding AI. The other AI in Fig. S1a can be derived in a similar manner using an arbitrary closed-form expression.

Extrapolating from the example, **without loss of generality, an Active Interconnection (AI)** relating *M* connecting REs with states 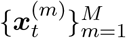 can be stated as:

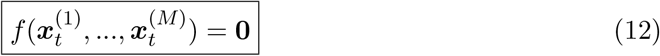

Here, 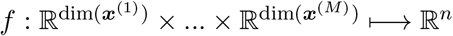 where *n* ∈ ℤ^+^ (positive integers).

Note, each state variable 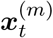 is associated with parameters that characterize its probability distribution. In the scope of above instantiation of RE (modified EKF), each state variable is associated with its mean and covariance. That is, 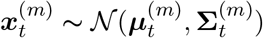, which is equivalent to 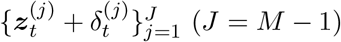, where 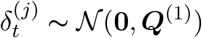.

#### S1.5 Combining REs and AIs

As it may be apparent by now, an AI (Eqn. 12) and an observation model of an RE (Eqn. 9) follow a similar form by design to enable tight interoperation between the two functional elements: An AI relates multiple REs, and each RE use their connecting AIs to perform inference. And the **ports** on REs serve to transmit information in the first case, and receive information in the second case. All put together, REs perform a joint estimation of several quantities/state in the context of the scaffold setup by the AIs.

This results in recovering complex structures in information while remaining robust to noise because AIs pass probabilistic information. This is crucial to perform Bayesian inference, such as in our work. That is, each RE uses the information obtained from other REs in a manner consistent with their probabilistic distribution. In the scope of this work, using the modified EKF mentioned above, this means each RE considers the uncertainty (covariance) of values received from other REs and weighs its own estimate according to the Bayes rule.

Please note, using the conventions we described above, sensory variables or values from other REs can be treated in the same way. That is, irrespective of the source of input (either from the external world, or from within a network), all connections to REs can in principle be treated as AIs. **However, for clarity of exposition when more than one RE is involved in a relation, we call this relation as AI (Eqn. 12), while when an RE has a relation only from sensory variables (characterized by an input-only *port*), we refer to this relation as an observation model (Eqn. 9)**.

### S2 Model Details for “Illusion 2: Silencing by Motion”

Here we will describe the model from Fig. 4a using the conventions mentioned in the previous section. In order to ease into details, we first describe the states (or quantities) involved in the model and explain the high-level relations those have to each other and external signals. We then setup AIs for relevant relations, and finally describe the REs that embody these characteristics.

#### S2.1 States and Relations

The motion RE block in Fig. 4a is comprised of a bank of REs one for every object (dot) in the stimulus. The state corresponding to an RE belonging to an object is given by:

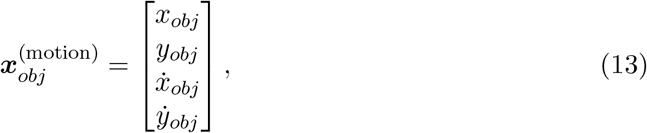

where *x*_*obj*_, *y*_*obj*_ are the x and y coordinates of the centers of the objects, in normalized units [0, 1]. And 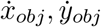 are the respective object velocities in normalized units*/s*.

In a similar manner, luminance RE block in Fig. 4a contains several REs, one for every object (dot) in the stimulus. The state corresponding to an object is given by:

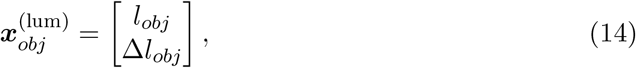

where *l*_*obj*_ denotes the luminance level of that object, in normalized units [−1, 1] (−1 = black, 0 = gray, 1 = white), and Δ*l*_*obj*_ is the change in luminance for that object between time steps, in the same units.

What signals and relations can be used to *infer* the states 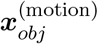 and 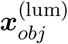?Refer-ring back to Fig. 4a, we see there are three sources of signal (depicted in the blue boxes), one AI between motion and luminance RE blocks, and AIs between several instances of object (dot) motion REs. We will first briefly describe each type of relation as points. In subsequent text, we provide more details for each of those relations.

##### Points

1. Motion of each object 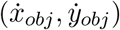 can be inferred from observations of positions of respective objects 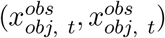. This corresponds to observation model 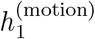.
2. The luminance of each object (*l*_*obj*_) may be recovered from “slow” luminance intensities on the “imaging surface” 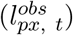, which corresponds to the observation model 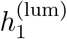.
3. “Fast” luminance changes 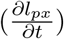 on the “imaging surface” behaves in a way consistent with motion 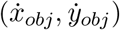 and luminance information for each object (*l*_*obj*_). This regularity can be used to supplement both luminance and motion perception. This is encoded with AI *f*_1_, and manifests as observation models 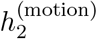 and 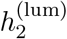.
4. Estimation of motion of each object can benefit from its neighbors. This is encoded with AI *f*_2_, and manifests as observation model 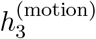.

#### S2.2 Active Interconnections

The following function addresses point 3 from above:

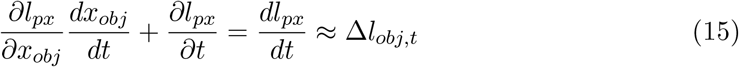

This is nothing but the total derivative of the spatiotemporal function *l*_*px*_(*x, t*) expressed in its partial derivatives. A similar form is used to recover optic flow under “brightness constancy” assumption [4], i.e., assuming 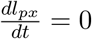 and 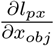 becomes akin to image spatial gradients, 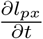 akin to temporal gradient, and 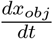 akin to optic flow vector.

Therefore, we can use Eqn. 15 and express it as an AI using our convention (Eqn. 12):

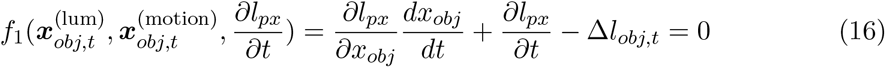

Point 4 can be expressed as the tendency for neighboring dots to “agree” with each other. If they agree, they must have same velocities. Therefore, the corresponding AI is straight-forward for objects in a neighborhood:

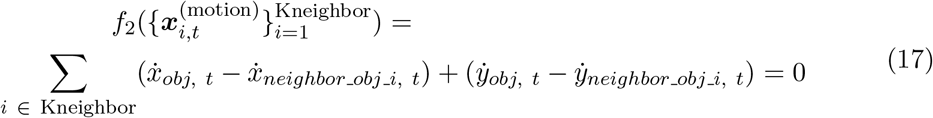

#### S2.3 Recursive Estimators

Now that we stated the different quantities (states) that are modeled in the system and how they can relate to each other, we can now detail the REs that perform the inference of those states from sensory signals and working with each other.

##### S2.3.1 Motion RE

**Process model** *g*^(**motion**)^ The RE for recovering motion per object can be specified with a constant velocity heuristic-based process model:

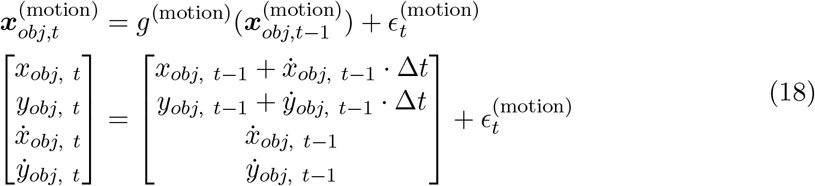

**Observation model** 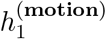 Following point 1 from above, an observation model is straight-forward:

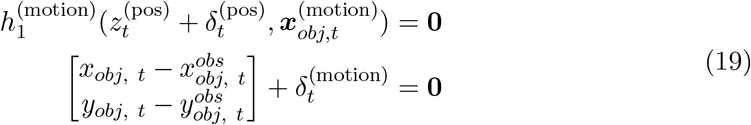

Note, the dot position as an observation is just a stand-in for second order motion perception [5], i.e., motion perception through object-level correspondences. There could be alternate motion estimation mechanisms, e.g., first-order motion perception similar to spatiotemporal energy [6], which possibly uses the “fast” path signal indirectly. Motion recovery in the human visual system may be complicated and multi-path. So, we simplify here for the sake of uncovering the interaction between motion and luminance perception through the lens of the phenomenon.

Also, this observation model does not employ outlier rejection.

**Observation model** 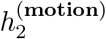 Motion can be recovered using the interaction between “fast” luminance changes and estimates of luminance as outlined in the AI given by Eqn. 16. The corresponding observation model is given by:

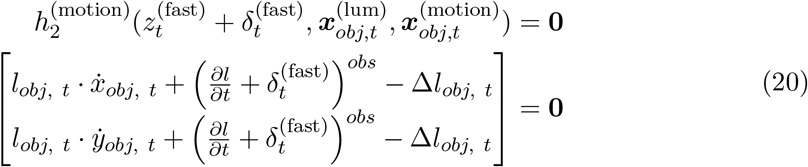

Here, (*l*_*obj*, *t*_, Δ*l*_*obj*, *t*_) are from luminance RE, and thus carry uncertainty information implicitly (it is not a simple additive term such as with 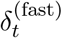, and this observation model employs outlier rejection threshold at a Mahalanobis distance of 1.

**Observation model** 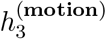 For improving collective estimation of motion, the observation model can be directly specified following the AI outlined in Eqn. 17:

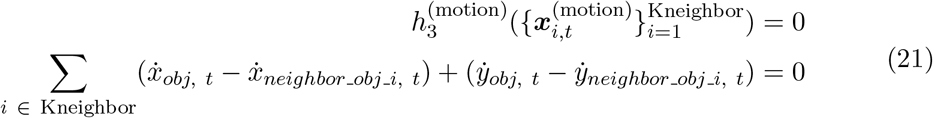

For this observation model, outlier rejection threshold is set at a Mahalanobis distance of 1*/*1000. Mechanistically, this means the “observations” from the neighbors are only effective when motion estimates of those neighbors are very close as well. The “closeness” is also proportional to this dot’s own confidence. That is, with more uncertainty, this dot becomes more susceptible to neighbor’s influence, whereas with low uncertainty, the likelihood of categorizing a neighbor’s motion as an outlier goes up.

##### S2.3.2 Luminance RE

**Process model** *g*^(**lum**)^ The process model for luminance recovery per object is specified as a constant velocity heuristic along with clipping:

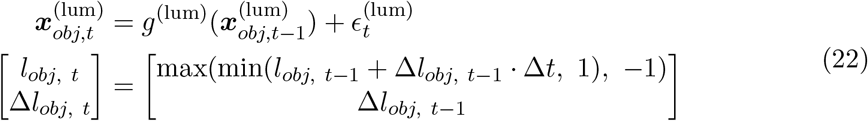

**Observation model** 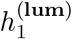 Following point 2 from above, the observation model pertaining to accumulating “slow” luminance values becomes

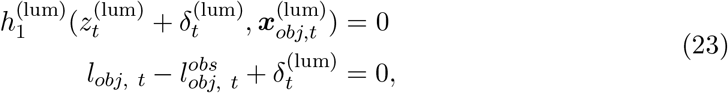

with no outlier rejection.

**Observation model** 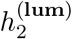 Similar to 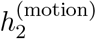, luminance can be recovered using the interaction between “fast” luminance changes and motion as outlined in the AI given by Eqn. 16. The corresponding observation model is given, which is identical to 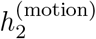except for the inputs is given by:

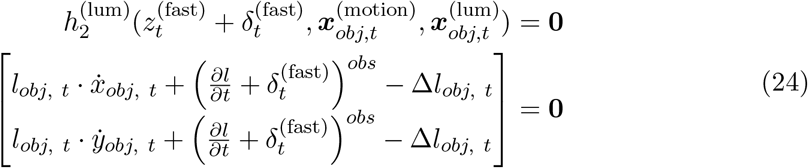

Here, 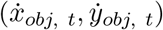 are from motion RE. The quantities carry uncertainty information implicitly. This observation model does not include outlier rejection.

### S3 Model Details for “Illusion 1: Fill-in Color After-Effect Illusion”

In this section, we will describe the model from Fig. 2a.

#### S3.1 States and Relations

The shape RE block in Fig. 2a consists of multiple REs. The number of REs is dynamically adjusted depending on the existence of star-shaped objects in the input stream. In the input stream, when an unrecognized shape (identified by location and orientation) appears, a new RE is created to track the instance of that shape. The state of each RE here is given by:

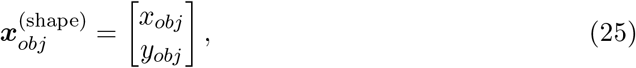

where *x*_*obj*_, *y*_*obj*_ are the x and y coordinates of the centers of the star-shaped objects, in units of pixels. In addition, each RE carries an auxiliary information about the type of star-shaped object 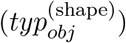: either axis-aligned star (similar to the 2nd input frame in Fig. 2a) or a diagonally-aligned star (similar to the 3rd input frame in Fig. 2a).

The color RE block in Fig. 2a consists of a 2D-bank of REs, one for every pixel in the input. Each RE here has the following state:

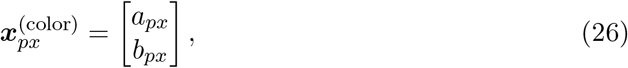

where *a*_*px*_, *b*_*px*_ correspond to Cartesian coordinate equivalent of hue and saturation (which are normally represented in polar coordinates) of the tracked pixel.

The states 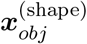 and 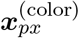 can be independently inferred from the input signals (depicted in the blue boxes of Fig. 2a,) translating to two relations between states and available signals. But additionally, the phenomenon indicates some interaction between shape and color processing, which is the third relation. First, let us briefly describe the relations as points. In subsequent text, we give more details for those relations.

##### Points

1. The existence and position of a shape (*x*_*obj*_, *y*_*obj*_) can be inferred directly from detections of corresponding shapes. This is given by the observation model 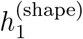.
2. The color of each pixel *a*_*px*_, *b*_*px*_ can be inferred from hue and saturation observed at the corresponding pixel in input. This corresponds to observation model 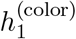.
3. To characterize interaction between shape and color, we can follow a naive assumption that correlations between shapes and colors in natural images statistics is higher than random. This statistical regularity can be used to improve mutual confidence of shape and color estimation. At the time of implementation of this model, an AI had a slightly different convention than the one mentioned in Sec. S1. So, we provide a more direct description of encoding of this statistical regularity.

#### S3.2 Recursive Estimators

##### S3.2.1 Shape RE

**Process model** *g*^(**shape**)^ This RE uses a trivial process model:

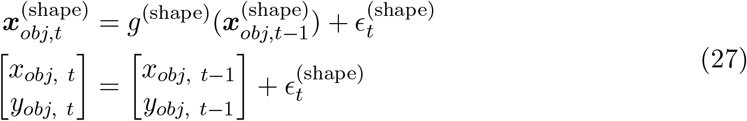

This introduces a “zeroth-order sluggishness” to the RE, where during inference, the current estimate of the state 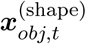 would not “like” to stray too far away from the the previous estimate 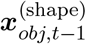.

**Observation model** 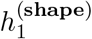 Following point1 from above, we can specify a straight-forward observation model:

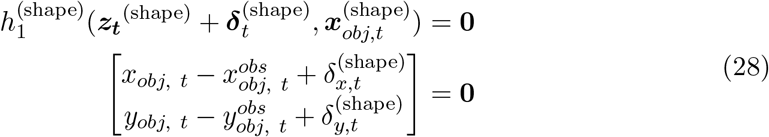

Here, the observations 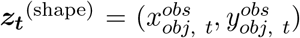 come from a shape detection algorithm that extracts star-like shapes from the brightness input stream. The observation noise 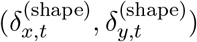 is scaled according to the “distance” between expected star-shape and the detected star-shape. That is, when star-shape being tracked in this RE is an axis-aligned star, the distance of a measured shape of the same orientation is 0, and the distance grows for deviations from this orientation with the maximum at 22.5 deg. Higher orientations will result in the distance being closer to diagonally-aligned star (which is a 45 deg rotation of the axis-aligned star), and thus categorized as a star of another type. The distance metric is obtained from normalized central moments of the contour of the shape.

##### S3.2.2 Color RE

**Process model** *g*^(**color**)^ The color RE for each pixel also uses a trivial process model:

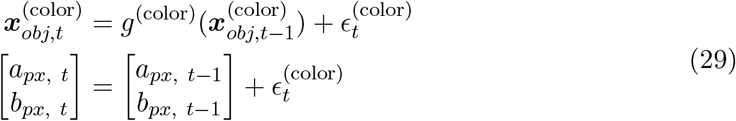

The reasoning behind this process model is the same as the that of *g*^(shape)^.

**Observation model** 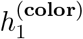 In accordance to point 2, the observation model here is given by:

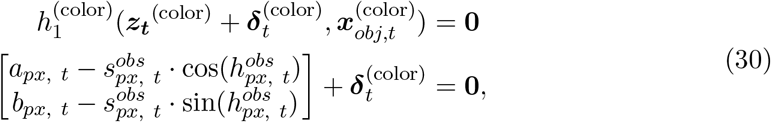

The observation noise 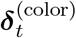 is modulated depending on how much “color” exists in the corresponding pixel.

#### S3.3 Interaction Between Shape and Color

The interaction between shape and color processing is implemented as a direct modulation of the uncertainty parameters of their respective states, i.e., 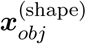 and 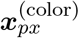.

From the perspective of the shape RE, the color RE (with all its bank of REs combined) represents an image stream similar to the brightness input stream, but in a different representation. This can be used as an alternate source of input for shape detection. Thus, whenever an already existing shape RE detects the same shape from color information, the corresponding shape RE’s uncertainty is reduced:

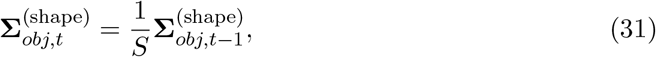

where *S >* 1 and 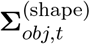 is the uncertainty 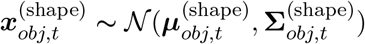.

At the same time, the color REs that remain within the confines of the shape are adjusted to have a lower uncertainty:

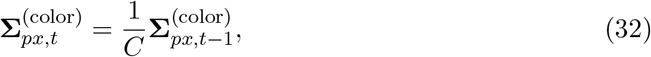

where *C >* 1 and 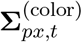 is the uncertainty of 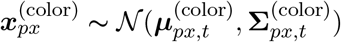.

This interaction introduces an unexpected but beneficial correlation between shape and color processing: each individual color RE within the block is not setup to *directly* interact with each other—the observations of color only affect the respective pixels. But through this interaction between shape and color, color REs within a particular shape receive information indirectly through a shape RE.

**Figure S1:**
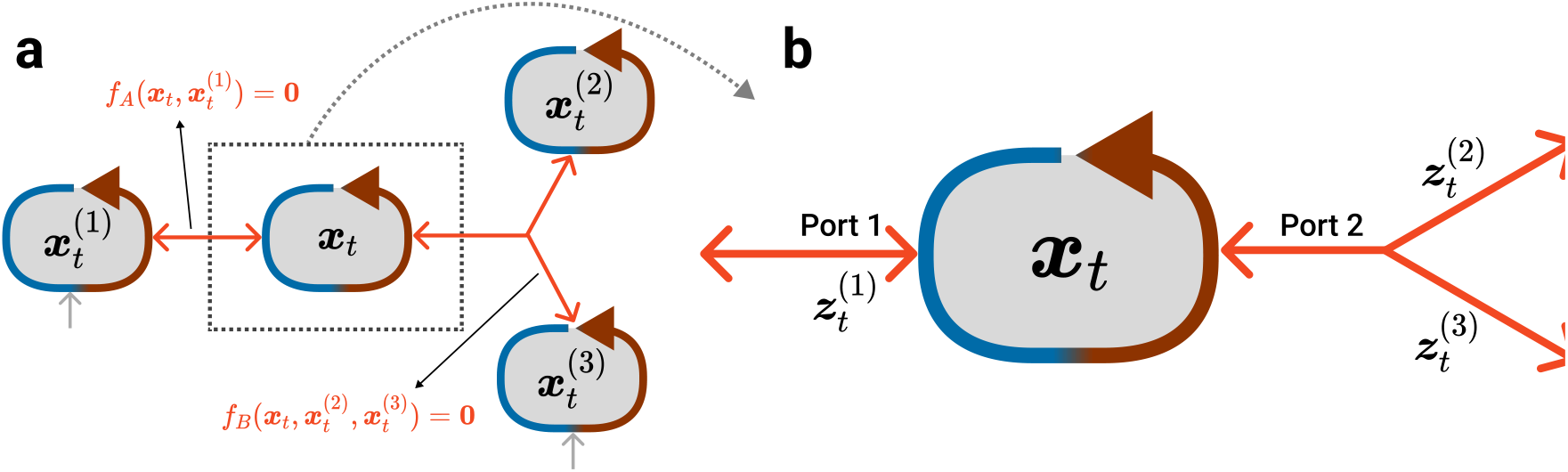
A toy example to illustrate functional elements of AICON. (a) Four Recursive Estimators (RE) are interconnected through two Active Interconnections (AI). (b) A zoomed in version of the same topology is provided, to illustrate reception of information at ports of REs. Please refer to text in SI Sec. S1 for more details.

**Figure S2:**
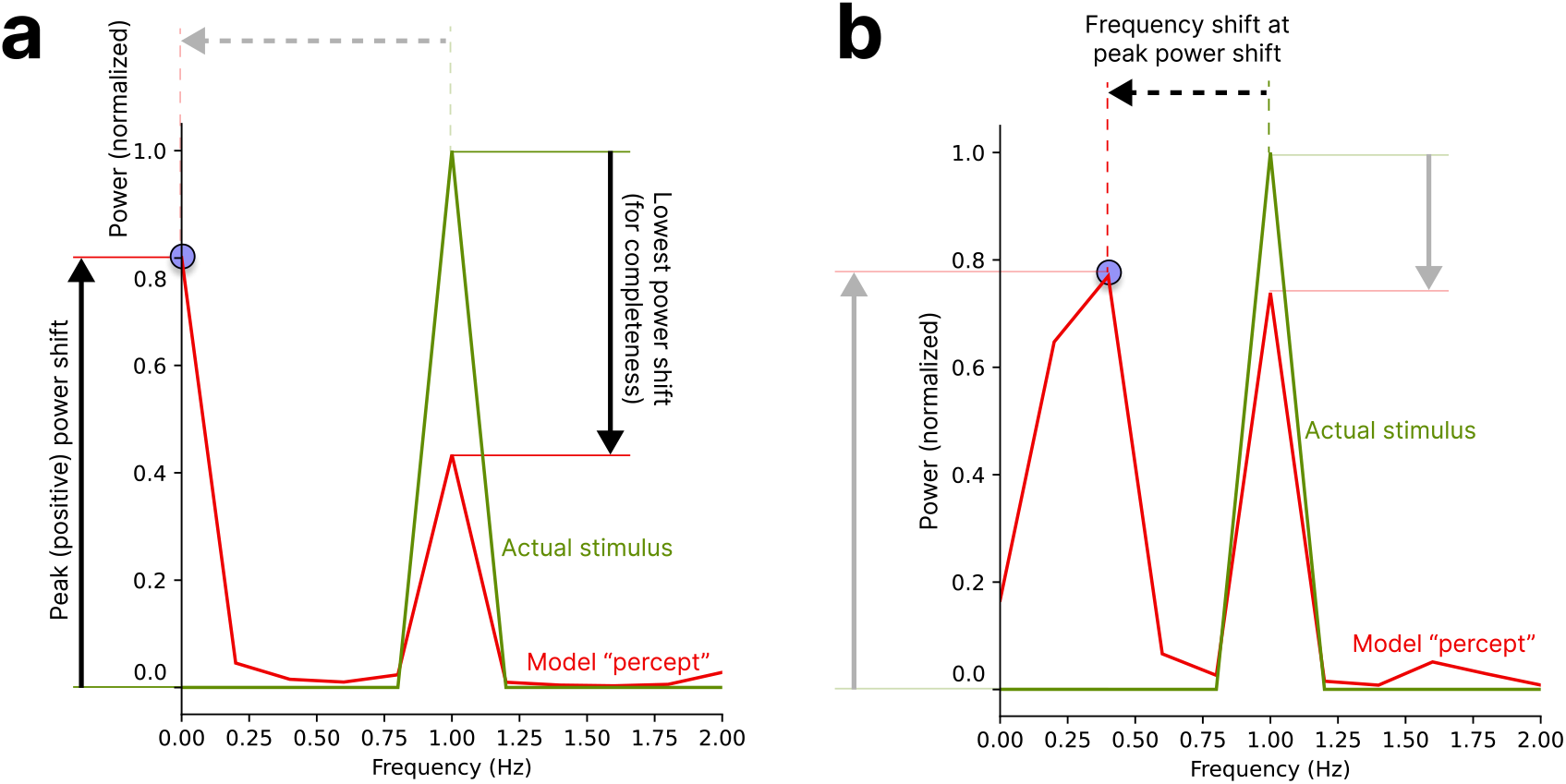
Peak power shift serves as a proxy to measure silencing. Here, the two plots correspond to model subjected to two different types of input stimuli. The plots depict the combined power spectrum of each dot in the stimuli (normalized by the number of dots). Therefore, the “actual stimulus” (shown in green) has a peak power of 1.0, at the stimuli frequency of 1 Hz. “Model percept” (shown in red) shows power distributed across different frequencies. The blue dots in both plots show the power shift-frequency pair considered to evaluate silencing. Peak power shift gives a continuous (a) Illustration of power shift metrics: At each frequency, the relative shift of power between “actual stimulus” and “model percept” is first obtained. The highest positive shift in power (shown as solid arrow) is dubbed “peak power shift”. For completeness, this diagram also shows lowest power shift that occurs at the actual stimuli frequency—this is not considered to measure degree of silencing. (b) Depiction of the frequency shift at the peak power shift: The “model percept” exhibits a significant shift in power at a frequency different from the “actual stimulus” frequency of 1 Hz. This can also serve as a measure of silencing if dominant frequency, such as in identifying musical notes, is indicative of the perceived frequency. We describe frequency shift here only for completeness. Frequency shift in practice has lesser granularity than power shift as a consequence of using Discrete Fourier Transform (DFT) that limits the number of frequencies for comparison. In our experiments, we found that peak power shift was sufficient to qualitatively explain the trends observed in humans, but deriving a metric that uses both peak power shift and frequency shift might be prudent for comparing further psychophysical experiments.

**Figure S3:**
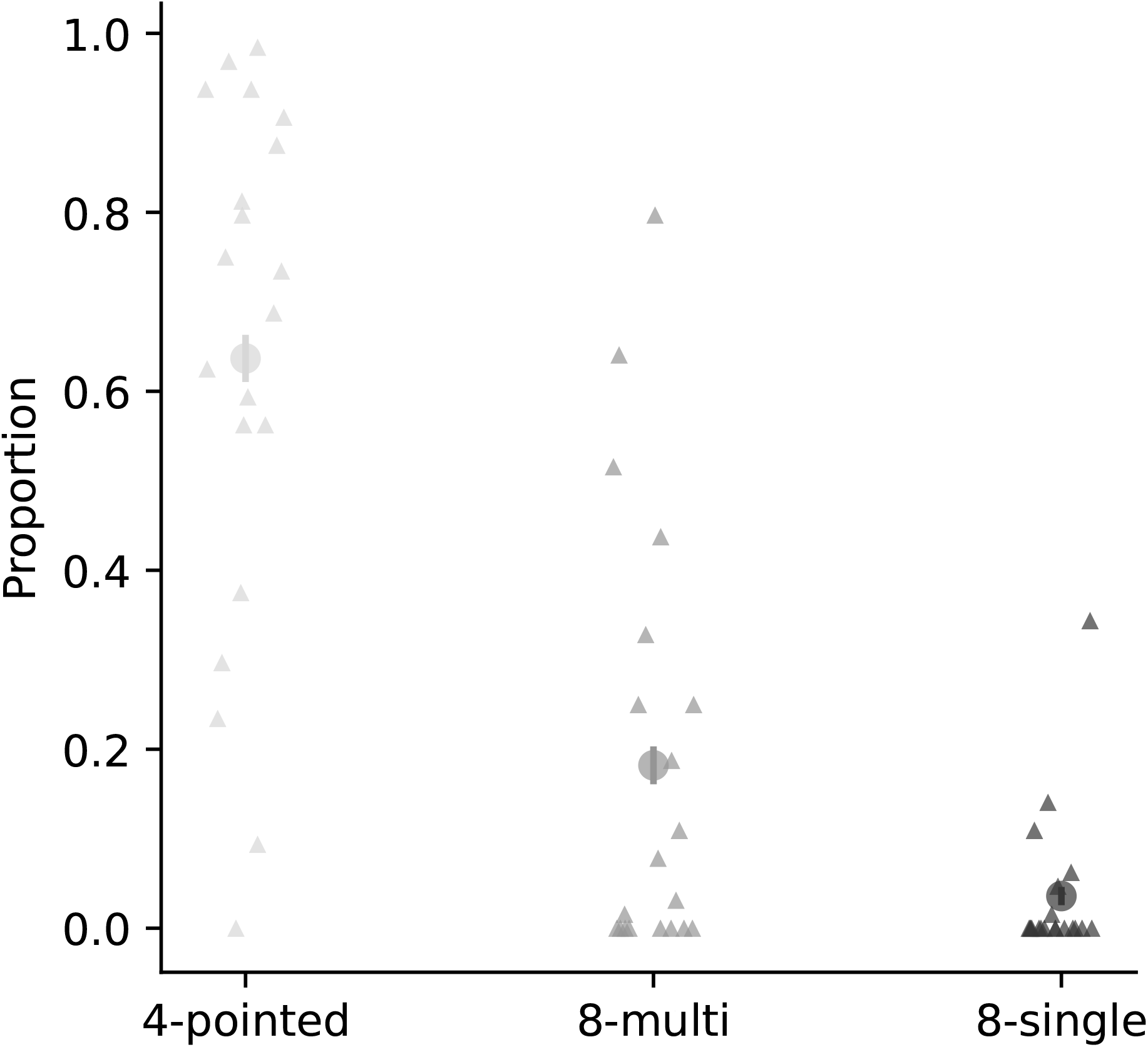
Proportions of perceiving an after-image per response for Experiment 1B. Each triangle represent the proportion of responses per after-image response option (4-pointed star: light gray; 8-pointed multi-colored: medium gray; 8-pointed single-colored: dark gray) for one participant. The proportion mean is indicated by the larger circle and the binomial confidence interval by the vertical line.

### Other Supplementary Material

- **Movie S1**. “Illusion 1: Fill-in Color After-Effect Illusion”: Demo movie
- **Movie S2**. “Illusion 1: Fill-in Color After-Effect Illusion”: Example stimuli used in experiments. Condition: No rotation, no size change
- **Movie S3**. “Illusion 1: Fill-in Color After-Effect Illusion”: Example stimuli used in experiments. Condition: 7 deg rotation, no size change
- **Movie S4**. “Illusion 1: Fill-in Color After-Effect Illusion”: Example stimuli used in experiments. Condition: No rotation, 120% size change
- **Movie S5**. “Illusion 2: Silencing by Motion”: Demo movie
- **Movie S6**. “Illusion 2: Silencing by Motion”: Example stimuli used in Exp. 2A (incoherent motion)
- **Movie S7**. “Illusion 2: Silencing by Motion”: Example stimuli used in Exp. 2B (only random walk motion)
- **Movie S8**. “Illusion 2: Silencing by Motion”: Example stimuli used in Exp. 2B (random walk motion & coherent motion)

## Notes

### Competing Interest Statement

The authors have declared no competing interest.

### Summary of Updates

"Unknown" term added to title; Acknowledgments updated; Continued page numbering into supporting information

